# PETScan: Score-Based Genome-Wide Association Analysis of RNA-Seq and ATAC-Seq Data

**DOI:** 10.1101/2025.04.10.648278

**Authors:** Yajing Hao, Tal Kafri, Fei Zou

## Abstract

High-dimensional sequencing data, such as RNA-Seq for gene expression and ATAC-Seq for chromatin accessibility, are widely used in studying systems biology. Accessible chromatin allows transcription factors and regulatory elements to bind to DNA, thereby regulating transcription through the activation or repression of target genes. The association analysis of RNA-Seq and ATAC-Seq data provides insights into gene regulatory mechanisms. Most existing analytic tools exclusively focus on cis-associations, despite regulatory elements being able to physically interact with distant target genes. Furthermore, conventional approaches often utilize Pearson or Spearman correlations, which ignore the count-based nature of RNA-Seq data.

To address these limitations, we introduce *PETScan*, a computationally efficient genome-wide ***PE***ak-***T***ranscript ***Sc***ore-based association ***an***alysis, utilizing negative binomial models to better accommodate RNA-Seq data. We leverage score tests and matrix calculations for improved computational efficiency, and combine an empirical permutation method with genomic control to ensure valid p-value calculations in studies with limited sample sizes. In real-world datasets, *PETScan* achieved three orders of magnitude faster than Wald tests, while identifying similar significant gene-peak pairs. The *PETScan* R package is available on GitHub at https://github.com/yajing-hao/PETScan.

## 1 Introduction

Transcriptomics data analysis investigates RNA transcripts produced by the genome under specific biological conditions or in particular cell types, which plays a significant role in understanding complex traits and diseases [Casamassimi et al., 2017]. Although every cell in the human body shares an identical set of genes, distinct gene expression patterns define the specialized functions of different cell types both physiologically and pathologically [Ralston and Shaw, 2008, Zeng, 2022]. These patterns can be regulated by epigenetic mechanisms, such as DNA methylation [Dhar et al., 2021] and histone modifications [Gagnidze and Pfaff, 2022], which regulate gene expression through modifications and structural changes to DNA without altering the underlying DNA sequence. Additionally, changes to chromatin accessibility represent another important epigenetic mechanism that regulates gene expression [Tsompana and Buck, 2014]. Structurally, a chromatin region can be densely packed (heterochromatin), leading to gene silencing, while loosely packed chromatin (euchromatin) tends to be transcriptionally active [Huisinga et al., 2006]. The dynamic nature of chromatin accessibility plays a crucial role in gene regulation by allowing transcription factors (TFs) and other regulatory elements to bind to DNA, thereby regulating transcription through the activation or repression of target genes [Klemm et al., 2019].

High-throughput sequencing technologies, such as RNA sequencing (RNA-Seq) [Nagalakshmi et al., 2008] and Assay for Transposase-Accessible Chromatin sequencing (ATAC-Seq) [Buenrostro et al., 2013], enable the study of gene expression and chromatin accessibility, respectively. When RNA-Seq and ATAC-Seq data are jointly collected from the same samples or cells, systematic investigations of the relationship between gene expression and chromatin accessibility can be conducted to identify chromatin regions that regulate gene expression and reveal specifically impacted genes. Such analysis sheds light on the regulatory mechanisms underlying chromatin accessibility [Starks et al., 2019]. Understanding these gene regulatory mechanisms is crucial for interpreting transcriptomic differences across various biological conditions, unraveling the progression of diseases [Roadmap Epigenomics Consortium et al., 2015], and developing effective therapeutic strategies [Dai et al., 2024]. For example, research has identified disease-specific genes and uncovered their regulatory mechanisms underlying chromatin accessibility in diabetes [Ackermann et al., 2016], hepatocellular carcinoma [Yang et al., 2021, Bai et al., 2024], cellular senescence [Song et al., 2022, Ding et al., 2023], and asthma [Zhang et al., 2023]. These studies provide insights into how genetic and epigenetic factors contribute to disease development and progression.

To identify associations between gene expression and chromatin accessibility, most existing analyses rely on Pearson correlation [Zhu et al., 2019, Chen et al., 2019, Stuart et al., 2021] or more robust Spearman correlation [Ma et al., 2020, Kartha et al., 2022, Park and Rhee, 2024]. For instances, cisDynet calculates Pearson correlation between gene expression and peak accessibility [Zhu et al., 2023], while TRIPOD detects peak-TF-gene trio regulatory relationships using Spearman correlation [Jiang et al., 2022]. More advanced statistical methods include partial least-squares regression [Staitieh et al., 2023], LASSO (least absolute shrinkage and selection operator) [Cao et al., 2018], adaptive elastic-net regression [Ledru et al., 2024], and a tree-based non-linear regression method called gradient-boosting machine [Bravo González-Blas et al., 2023]. All these methods ignore the count-based nature of RNA-Seq data, which can result in suboptimal performance and biased results, necessitating specialized statistical methods that appropriately handle RNA-Seq data.

Furthermore, current association analysis of RNA-Seq and ATAC-Seq data primarily focuses on identifying cis-associations in cis-regulatory regions [Frankel, 2012, Klemm et al., 2019]. For instance, cisDynet evaluates the association between a gene’s transcriptomic expression and peak accessibility within 500 kb upstream and downstream of its transcription start site (TSS) [Zhu et al., 2023]. However, research has shown that gene regulatory elements bound to accessible chromatin can physically interact with distant target genes through chromatin looping [Kadauke and Blobel, 2009], thereby modulating genes located far along the same chromosome or even on different chromosomes [Miele and Dekker, 2008, Dean, 2011, Dekker and Misteli, 2015, Van Den Heuvel et al., 2015, Maass et al., 2019]. A recent study has generated a comprehensive meta-Hi-C chromatin contact map, providing both functional cis- and trans-chromosomal interactions [Lohia et al., 2022].

To fully understand how chromatin accessibility regulates gene expression, a holistic approach that considers both cis- and trans-associations, or genome-wide association analysis, is necessary. However, RNA-Seq data typically includes expression profiles for over 20,000 genes, while ATAC-Seq data contains peak measurements from approximately 100,000 chromatin regions. This results in roughly 20,000 × 100,000, or 2 billion, paired association analyses, requiring a computationally efficient method that is currently unavailable.

To address these issues, we introduce *PETScan*, a computationally efficient genome-wide ***PE***ak-***T*** ranscript ***Sc***ore-based association ***an***alysis. The algorithm *PETScan* employs negative binomial models to respect the count nature of RNA-Seq data with potential overdispersion [Robinson et al., 2010, Di et al., 2011, Love et al., 2014]. To mitigate the computational challenge, *PETScan* adopts two strategies to ensure its practical usage. First, *PETScan* utilizes score tests that require only parameter estimates under the null hypothesis. This feature becomes particularly useful since, for a given gene, its restricted estimates under the null hypothesis remain the same across all tested ATAC-Seq peaks and need to be evaluated only once.

Second, to further enhance computational efficiency, PETScan mimics Matrix eQTL [Shabalin, 2012], an ultra-fast expression quantitative trait loci (eQTL) analysis tool. eQTL analysis also involves billions of paired association tests between millions of single-nucleotide polymorphisms (SNPs) and tens of thousands of genes across the entire genome. While Matrix eQTL converts traditional linear models into correlation coefficients for efficient large matrix operations, we combine score tests with ATAC-Seq data transformation to facilitate efficient matrix calculations.

Furthermore, in studies with small sample sizes, score tests may fail to converge to their theoretical asymptotic distributions. To better control the family-wise error rate or false discovery rate, *PETScan* incorporates an empirical permutation method inspired by genomic control [Devlin and Roeder, 1999], which has been successfully used for robust statistical inference in differential gene expression analysis with RNA-Seq data [Zou et al., 2014]. We demonstrate the effectiveness of *PETScan* using ENCODE human tissue-specific data and 10x Genomics embryonic mouse brain data. Computationally, *PETScan* is three orders of magnitude faster than the equivalent Wald tests.

## 2 Methods

Assume there are *N* paired samples of RNA-Seq and ATAC-Seq data with *G* genes and *P* peaks, respectively. Let *x*_1*i*_, *x*_2*i*_, …, *x*_(*k*−1)*i*_ represent a set of covariates in the *i*th sample (*i* = 1, …, *N*), such as the top principal components derived from the RNA-Seq data, which are commonly used to account for batch effects. For gene *g* (*g* = 1, …, *G*) and peak *p* (*p* = 1, …, *P*), let *y*_*gi*_ be the expression level of gene *g*, and *z*_*pi*_ denote the log-transformed chromatin accessibility for peak *p* in the *i*th sample. For simplicity of notation, we drop *g* and *p* out of the subscripts when describing the model for testing the association between the gene *g* and peak *p* pair in the following discussion, whenever there is no confusion.

To adjust for potential over-dispersion, we assume that *y*_*i*_ follows a negative binomial distribution with the density function *f* (*y*_*i*_; *µ*_*i*_, *ϕ*), where *µ*_*i*_ represents the mean and *ϕ* denotes the dispersion parameter with E(*Y*_*i*_) = *µ*_*i*_ and 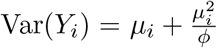. Specifically,

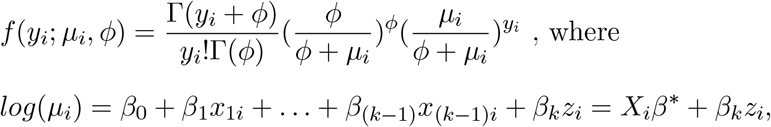

with, 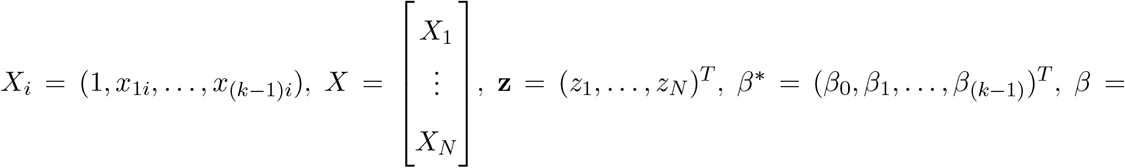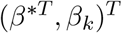 and *θ* = (*ϕ, β*^*T*^)^*T*^.

The log likelihood function of the data is therefore

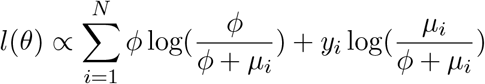

with the corresponding score function 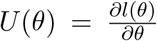 and the Fisher information matrix 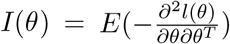. For the gene and peak pair, the primary interest is to test *H*_0_ : *β*_*k*_ = 0 versus *H*_*A*_ : *β*_*k*_ ≠ 0, which assesses whether the chromatin accessibility at the peak affects the expression of the gene. Let 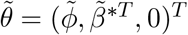 be the restricted maximum likelihood estimate of *θ* under the null hypothesis. Note that the parameter estimate 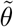 does not depend on the *z*_*pi*_. Therefore, for gene *g*, it remains consistent across all *P* peaks and only needs to be evaluated once.

For testing *H*_0_ : *β*_*k*_ = 0, *PETScan* uses the score test

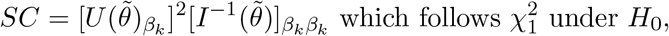

where 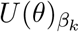 denotes the (*k* + 2)th element of *U* (*θ*), and 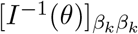 represents the (*k* + 2, *k* + 2)th entry of the inverse of *I*(*θ*), both corresponding to the parameter *β*_*k*_. Specifically, 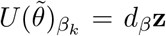, and 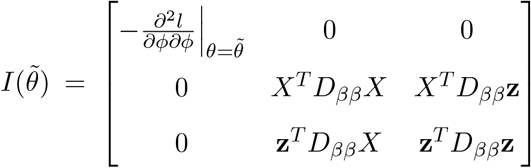, where 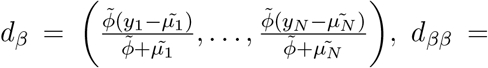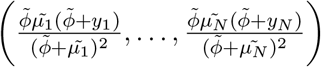, and *Dββ* = diag(*dββ*). Leveraging blockwise matrix inversion,

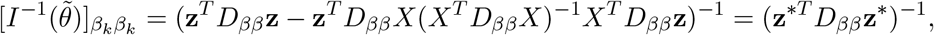

where **z**^*^ = [*I* − *X*(*X*^*T*^ *D*_*ββ*_*X*)^−1^*X*^*T*^ *D*_*ββ*_]**z**. After constructing **z**^*^ by applying a transformation to **z**, we can extract specific elements without computing the full inverse of a matrix.

Again, for gene *g, d*_*β*_ and *d*_*ββ*_ depend only on 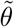 and remain consistent across all *P* peaks. Consequently, the score test for peak *p* can be extended easily to all peaks utilizing matrix calculations as described below: let **Z** = [**z**_1_, …, **z**_*P*_], **Z**^*^ = [*I* − *X*(*X*^*T*^ *D*_*ββ*_*X*)^−1^*X*^*T*^ *D*_*ββ*_]**Z**, then **SC** = (*SC*_1_, …, *SC*_*P*_) = (*d*_*β*_**Z**) ⊙ [*d*_*ββ*_(**Z** ⊙ **Z**)]^ℜ^ ⊙ (*d*_*β*_**Z**), where ⊙ represents the Hadamard (element-wise) product and *A*^R^ denotes the element-wise reciprocal of a vector or matrix *A*. Unlike the Wald and likelihood ratio tests, for a given gene, score tests require fitting the negative binomial model under the null hypothesis only once for all peaks, instead of *P* full models separately for each peak. This approach facilitates the proposed matrix operations to simultaneously calculate the score tests across all peaks, rather than repeated calculations for one peak at a time. This streamlines computations and significantly enhances computational efficiency.

For studies with finite samples, score tests may be artificially inflated, leading to inaccurate p-values derived from the asymptotic null distribution. To ensure robust inference, we combine an empirical permutation method with genomic control to mitigate concerns about asymptotic results. For gene *g*, let the original score test statistics across the *P* peaks be denoted as *SC* = (*SC*_1_, …, *SC*_*P*_). We then randomly sample the ATAC-Seq data without replacement for *L* times, and similarly denote the permuted score tests as 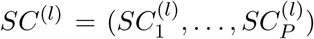,, where *l* = 1, …, *L*. Following the genomic control approach, we empirically estimate the inflation factor *λ* by 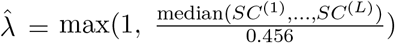 and rescale the original score test statistics *SC*_*p*_ to 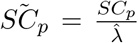, for *p* = 1, …, *P*, which are then compared to 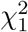 for p-value calculations.

## 3 Results

### 3.1 ENCODE Human Tissue-Specific Data

We applied *PETScan* to paired RNA-Seq and ATAC-Seq data from 46 healthy individuals across 15 human tissues, obtained from the Encyclopedia of DNA Elements (ENCODE) project [ENCODE Project Consortium, 2012], which were accessible at https://www.encodeproject.org/search/?type=Experiment. The RNA-Seq dataset contained 59,526 genes, and we applied filtering based on gene expression counts and selected variable genes, resulting in a final set of 11,637 genes. Similarly, the ATAC-Seq dataset included 805,169 peaks, which were filtered down to 132,670 peaks. We incorporated gender and the top two principal components derived from log-transformed gene expression data as covariates. P-values culminated in assessing associations across 1,548,656,910 gene-peak pairs. Leveraging parallel computation across 64 CPU cores (AMD EPYC 7313), the analysis was completed in approximately 40 minutes, significantly outperforming Wald tests, which required over 1,000 times longer. We identified 53,113 significant gene-peak pairs with Bonferroni-adjusted p-values below 0.05, suggesting a widespread impact of chromatin accessibility on gene expression.

For further analysis, we focused on the top 7,834 most significant gene-peak pairs with Bonferroni-adjusted p-values below 10^−5^, as visualized in Figure 1, which was adapted from [Sun, 2009]. To uncover over-represented pathways, we conducted Kyoto Encyclopedia of Genes and Genomes (KEGG) enrichment analysis on the genes associated with these top pairs using clusterProfiler [Yu et al., 2012]. Among enriched pathways, seven genes (*BHLHA15, FOXA2, FOXA3, HNF1A, HNF1B, HNF4A, HNF4G*) are involved in maturity onset diabetes of the young, six genes (*AMY1B, ATP2A3, KCNQ1, PLA2G10, RYR2, SLC4A4*) are related to pancreatic secretion, and four genes (*GLP1R, KCNMB1, RIMS2, RYR2*) are associated with insulin secretion, providing insight into the molecular mechanisms underlying pancreas-related diseases and metabolic disorders.

**Figure 1:**
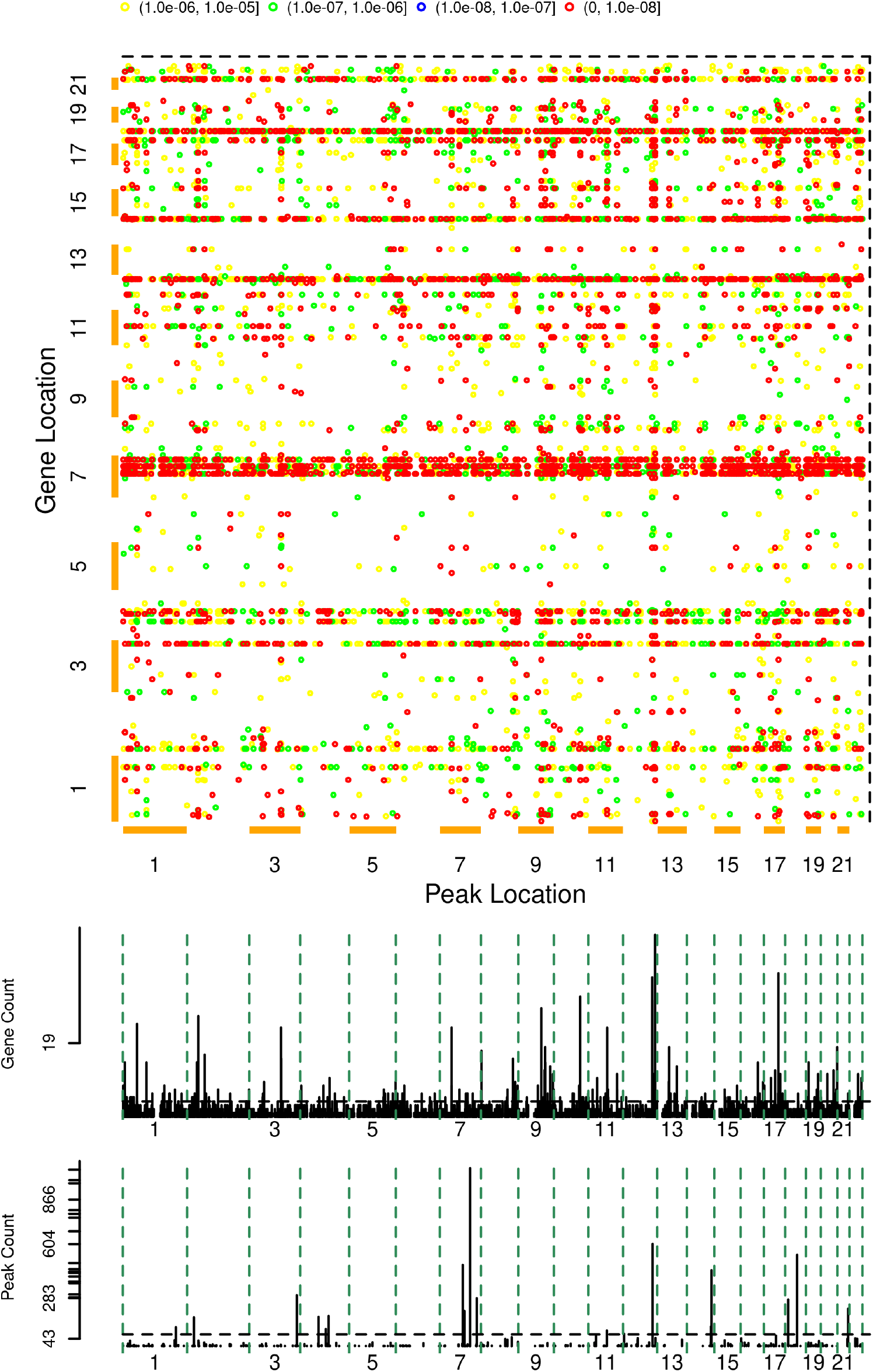
Significant gene-peak pairs across the genome from ENCODE human tissue-specific data are shown, with peak locations on the X-axis and gene locations on the Y-axis. Different levels of significance are indicated using different colors. The middle panel illustrates the number of genes associated with each peak, while the bottom panel displays the number of peaks linked to each gene.

One notable result involved an ATAC-Seq peak at chr12:124,871,497-124,872,709, which was associated with 47 genes across the genome, represented as a vertical line in Figure 1. Among these genes, *HNF1B, EGF*, and *SLC4A4* were linked to pathways for maturity onset diabetes of the young, pancreatic cancer, and pancreatic secretion, respectively. As shown in Figure 2, chromatin accessibility at this peak was negatively related to the expression of these associated genes. Notably, the pancreas exhibited lower chromatin accessibility at this peak yet higher gene expression, distinguishing it from other tissues. These findings highlighted the role of chromatin accessibility in tissue-specific gene regulation and shed light on the regulatory landscape of genes involved in pancreatic function and disease. For other associated genes, *BARX2* encoded the transcription factor Barx2, which regulated *GRHL2* ; in turn, the transcription factor Grhl2 may potentially target multiple downstream genes, including *BICDL2, CA12, CLDN4, INAVA, PKP3, RHOV*, and *TRPV6*, as per the Cistrome Data Browser, a comprehensive database for transcriptional and epigenetic regulation studies [Lambert et al., 2018,Mei et al., 2017,Zheng et al., 2019]. This regulatory cascade illustrated how chromatin accessibility at a specific peak could influence the coordinated expression of multiple genes across the genome. Such co-association suggested a shared regulatory mechanism, emphasizing the dynamic interplay between chromatin structure and gene regulation.

**Figure 2:**
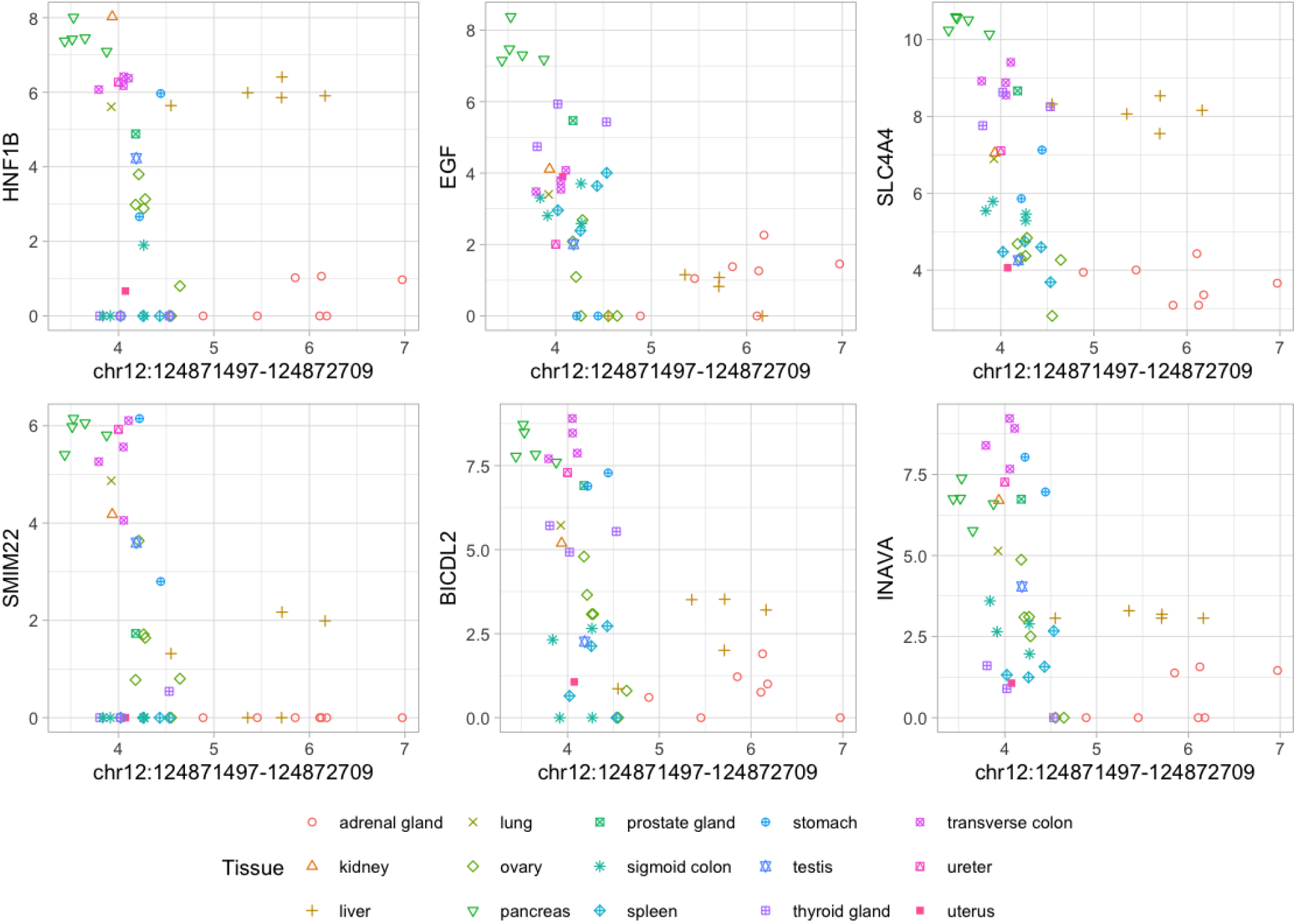
Scatter plots of chromatin accessibility at chr12:124,871,497-124,872,709 and the expression levels of its associated genes.

In addition, *PETScan* identified a few genes whose expression levels were associated with a large number of peaks, represented by horizontal lines in Figure 1. The most prominent was the *RPL32* pseudogene on chromosome 7, which originated from the retrotransposition of *RPL32* mRNA. This association primarily reflected tissue-specific gene expression and chromatin accessibility. Specifically, this gene exhibited higher expression in the pancreas while being lowly expressed in other tissues, while the chromatin accessibility of its associated peaks in the pancreas was either significantly higher or lower compared to other tissues (Figure 3). This finding was consistent with results from the GENCODE project, indicating that the transcription and regulation of pseudo-genes were tissue-specific [Pei et al., 2012]. A similar pattern was observed for the gene *ST8SIA5* on chromosome 18, which showed significantly higher expression in the adrenal gland, along with chromatin accessibility that was either much higher or lower in this tissue compared to others (Figure 3), suggesting a strong tissue-dependent regulatory mechanism. Existing research has shown that the expression of this protein-coding gene is linked to open chromatin regions, particularly in brain tissues [Ehrlich et al., 2024]. Our study did not include samples from brain tissues, but our analysis suggested that this highly expressed gene in the adrenal gland was also associated with open chromatin regions in the adrenal gland.

**Figure 3:**
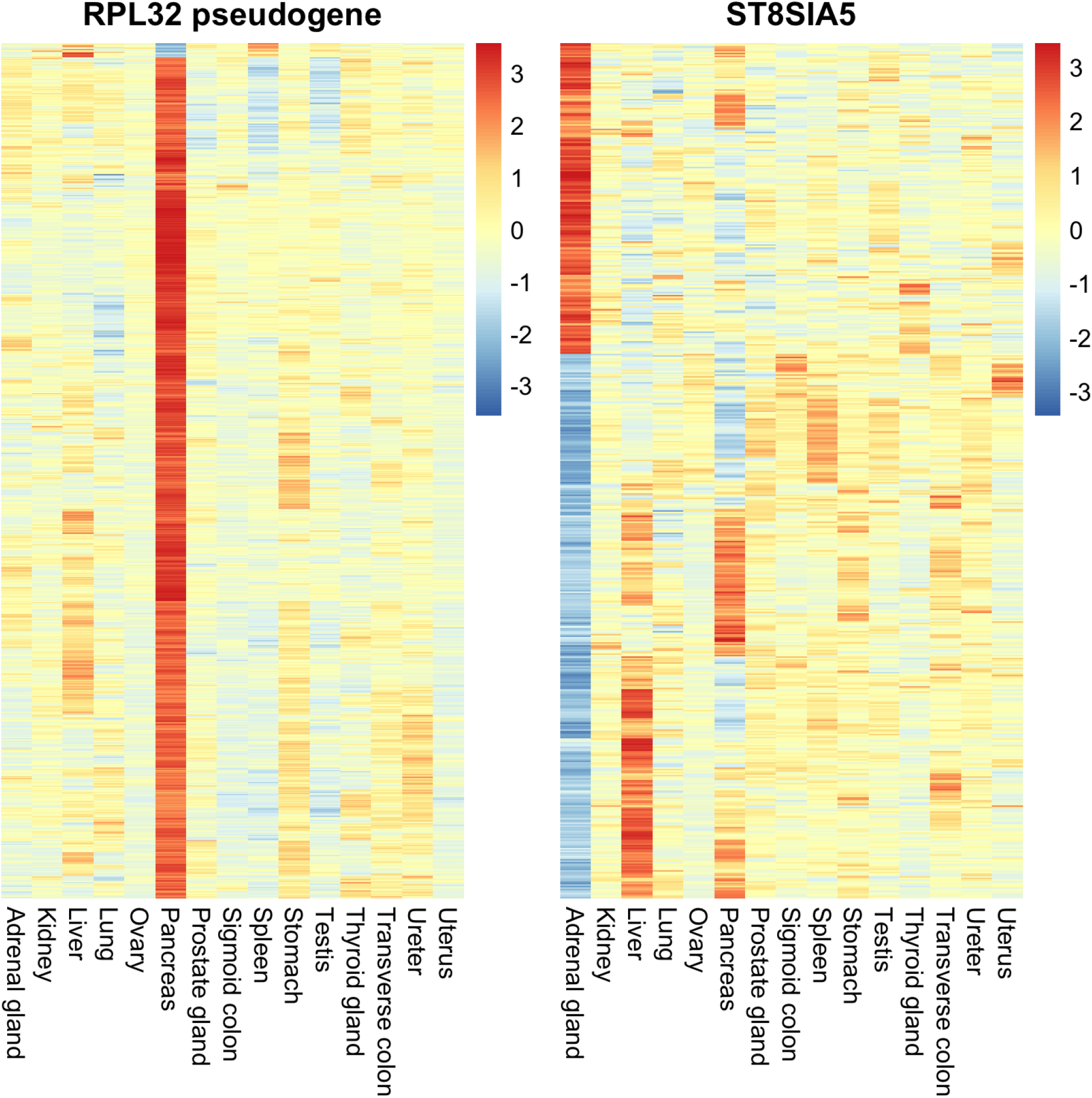
Heatmap of chromatin accessibility of associated peaks for the *RPL32* pseudogene (left) and *ST8SIA5* (right).

To evaluate our findings, we assessed whether significant gene-peak pairs were validated by previously reported cis-eQTLs from the Genotype-Tissue Expression (GTEx) analysis V8 release [Lonsdale et al., 2013], as cis-eQTLs data are more comprehensive than trans-eQTLs data, even though we investigated both cis- and trans-associations. Two gene-peak pairs overlapped with GTEx cis-eQTLs, where variants from significant variant-gene associations were located within peak regions, both exhibiting higher chromatin accessibility and higher gene expression in the pancreas, setting it apart from other tissues (Figure 4). For instance, variant rs4444903, situated within a peak spanning chr4:109,911,610-109,916,356, explained a fraction of the genetic variance in *EGF* expression. Overexpression of *EGF* and its receptor *EGFR* is frequently observed in pancreatic cancer, leading to uncontrolled cell proliferation and tumor progression, and is associated with poor prognosis [Heby et al., 2020]. This finding highlighted the tissue-specific accessibility of this region in modulating *EGF* expression, thereby contributing to pancreatic functionality and possibly to the pathological mechanisms underlying pancreatic cancer. Similarly, variant rs66817580 was linked to *FAM3B* expression, which was located within a peak at chr21:41,315,678-41,317,753. *FAM3B*, also known as *PANDER* (pancreatic-derived factor), is primarily expressed in the pancreas, particularly within beta cells. This gene is instrumental in the regulation of glucose metabolism and insulin secretion, and influences beta cell functionality and survival, making it relevant to diabetes and pancreatic disorders [Dong et al., 2023]. Further investigation into this regulatory mechanism could provide insights into beta cell dysfunction in diabetes and uncover potential therapeutic targets. *PETScan* demonstrated its effectiveness in uncovering gene regulatory mechanisms, particularly pancreatic-specific associations between chromatin accessibility and targeted gene expression.

**Figure 4:**
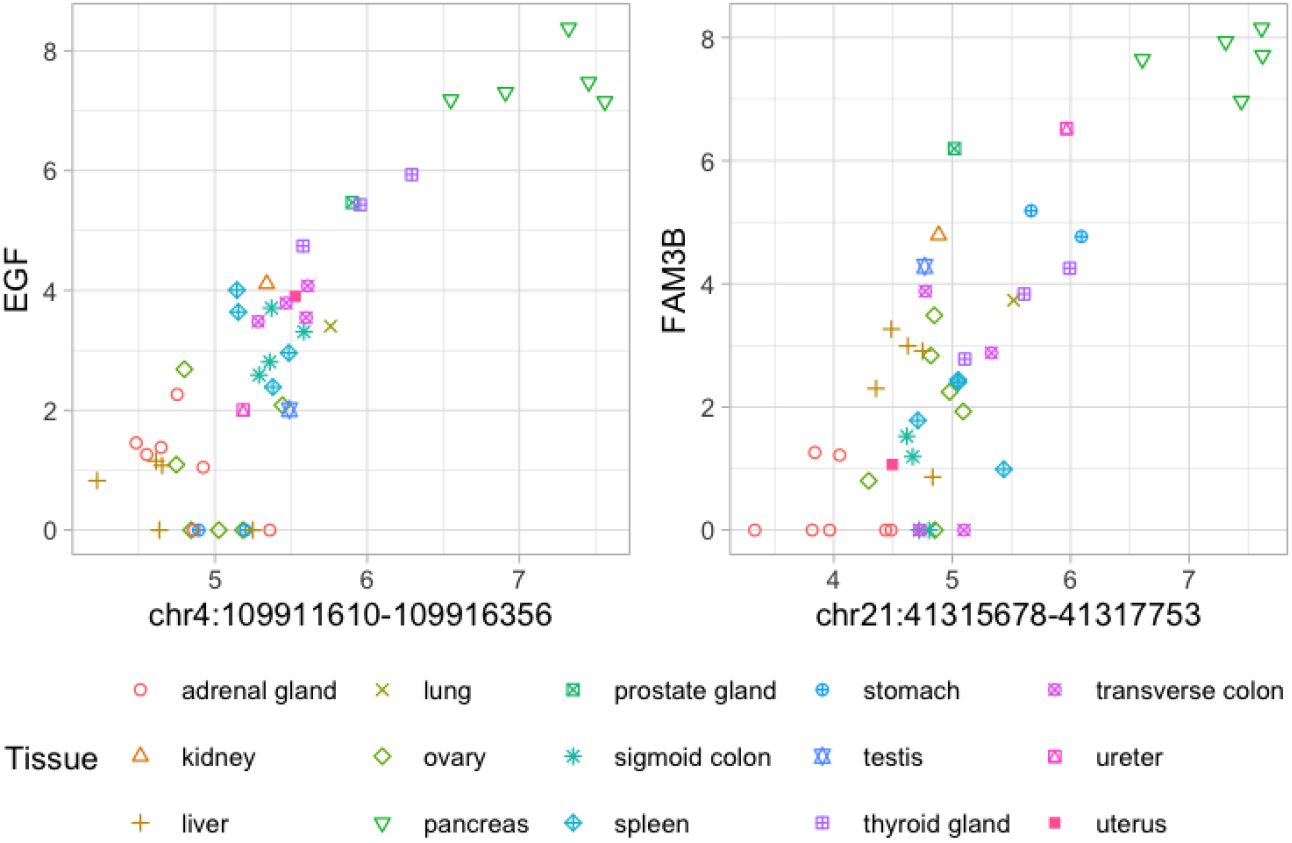
Scatter plots of significant gene-peak pairs from ENCODE human tissue-specific data validated by GTEx cis-eQTLs.

### 3.2 10x Genomics Embryonic Mouse Brain Data

We next conducted *PETScan* on publicly available 10x Genomics embryonic mouse brain data, which included 4,881 cells from the fresh cortex, hippocampus, and ventricular zone of the embryonic mouse brain at day 18 [10x Genomics, 2021]. The scRNA-Seq and scATAC-Seq data were processed following the TRIPOD vignette [Jiang et al., 2022], with cell-type labels transferred from an independent scRNA-Seq reference [La Manno et al., 2021] using SAVERCAT [Huang et al., 2020]. We retained 3,851 cells with consistent transferred labels from four major cell types: neuroblast, forebrain GABAergic neuron, cortical or hippocampal glutamatergic neuron, and glioblast. To address the sparsity issue in single-cell data, we partitioned cells into 84 metacells by clustering based on their scRNA-Seq profiles and assigned cell types to metacells according to the majority cell types within each cluster.

After filtering, we retained 5,935 genes and 42,048 peaks. Using parallel computation across 64 CPU cores, we analyzed 249,554,880 gene-peak pairs in just 7 minutes, adjusting for the first two principal components of the log-transformed gene expression data. *PETScan* demonstrated computational efficiency that was three orders of magnitude faster than Wald tests. We identified 26,991 significant gene-peak pairs with Bonferroni-adjusted p-values below 0.05, providing insights into the regulatory landscape of the embryonic mouse brain.

For further analysis, we focused on the top 6,858 significant gene-peak pairs with Bonferroni-adjusted p-values below 10^−5^, as illustrated in Figure 5. We performed Gene Ontology (GO) and KEGG enrichment analyses on genes from these pairs to identify over-represented GO terms and KEGG pathways using clusterProfiler [Yu et al., 2012]. The top enriched GO terms included forebrain development, telencephalon development, regulation of neurogenesis, gliogenesis, central nervous system neuron differentiation, and neuron projection guidance. These terms highlighted key processes involved in the development and specialization of the nervous system, particularly regarding the formation of the brain and its functional components. These findings aligned with the fact that embryonic mouse brain cells were collected during the transition phase between neurogenesis and gliogenesis, further supporting the validity of our results. The top enriched KEGG pathways grouped into two main categories: (1) nervous system: including glutamatergic synapse, GABAergic synapse, and synaptic vesicle cycle, all critical for neurotransmission and communication between neurons; (2) signal transduction: comprising calcium signaling pathway and neuroactive ligand-receptor interaction, which played essential roles in regulating cellular responses to external stimuli and modulating various physiological processes. These results were consistent with the characteristics of glutamatergic and GABAergic neurons included in embryonic mouse brain cells.

**Figure 5:**
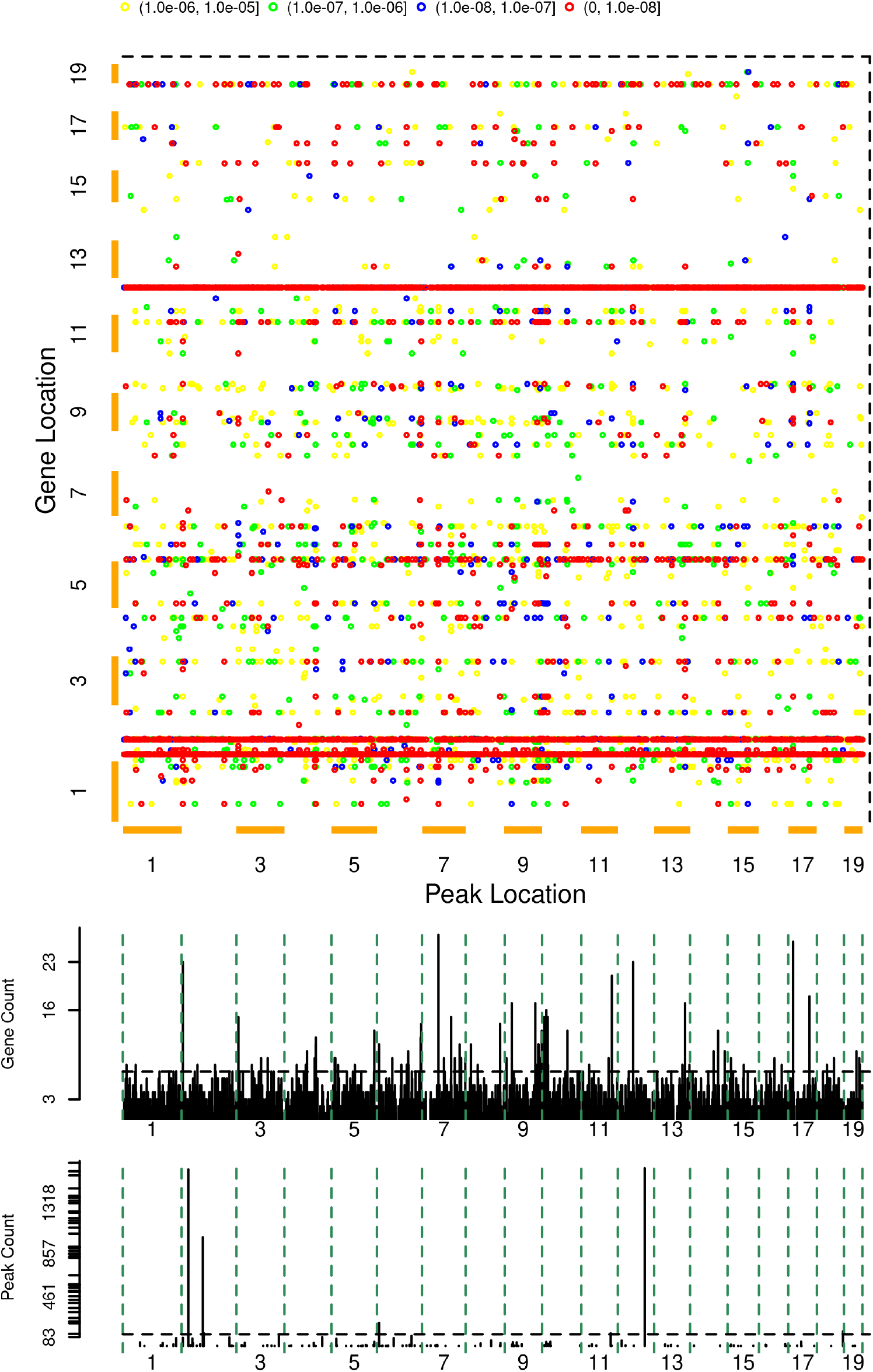
Significant gene-peak pairs across the genome from 10x Genomics embryonic mouse brain data are shown, with peak locations on the X-axis and gene locations on the Y-axis. Different levels of significance are indicated using different colors. The middle panel illustrates the number of genes associated with each peak, while the bottom panel displays the number of peaks linked to each gene.

Furthermore, an ATAC-Seq peak at chr7:54,633,595-54,635,192 was associated with 27 genes across the genome. Notably, open chromatin at this peak was linked to lower expression of *Nrxn3, Gad1*, and *Gad2* (Figure 6), all of which were involved in the GABAergic synapse. Conversely, it was correlated with higher expression of *Neurod6, Zbtb18, Nfix, Lhx2*, and *Id2* (Figure 7), all of which were essential for telencephalon development and forebrain development. The upregu-lated or downregulated genes exhibited functional coherence, indicating that changes in chromatin accessibility may co-regulate genes involved in related biological processes.

**Figure 6:**
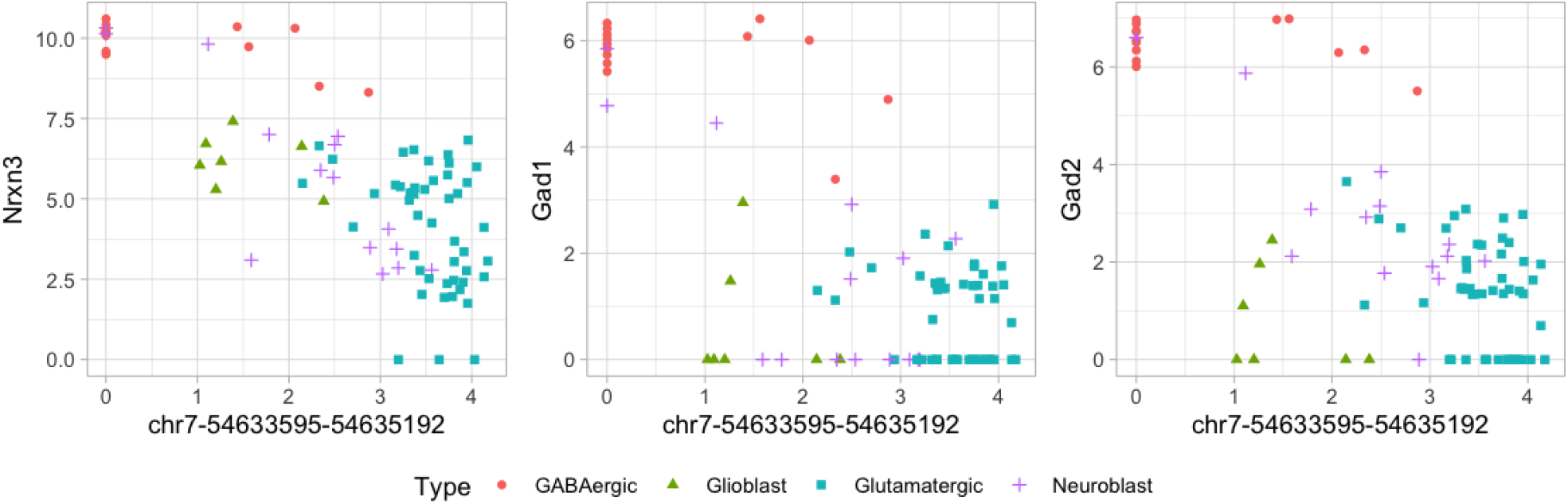
Scatter plots of chromatin accessibility at chr7:54,633,595-54,635,192 and the expression levels of its negatively associated genes.

**Figure 7:**
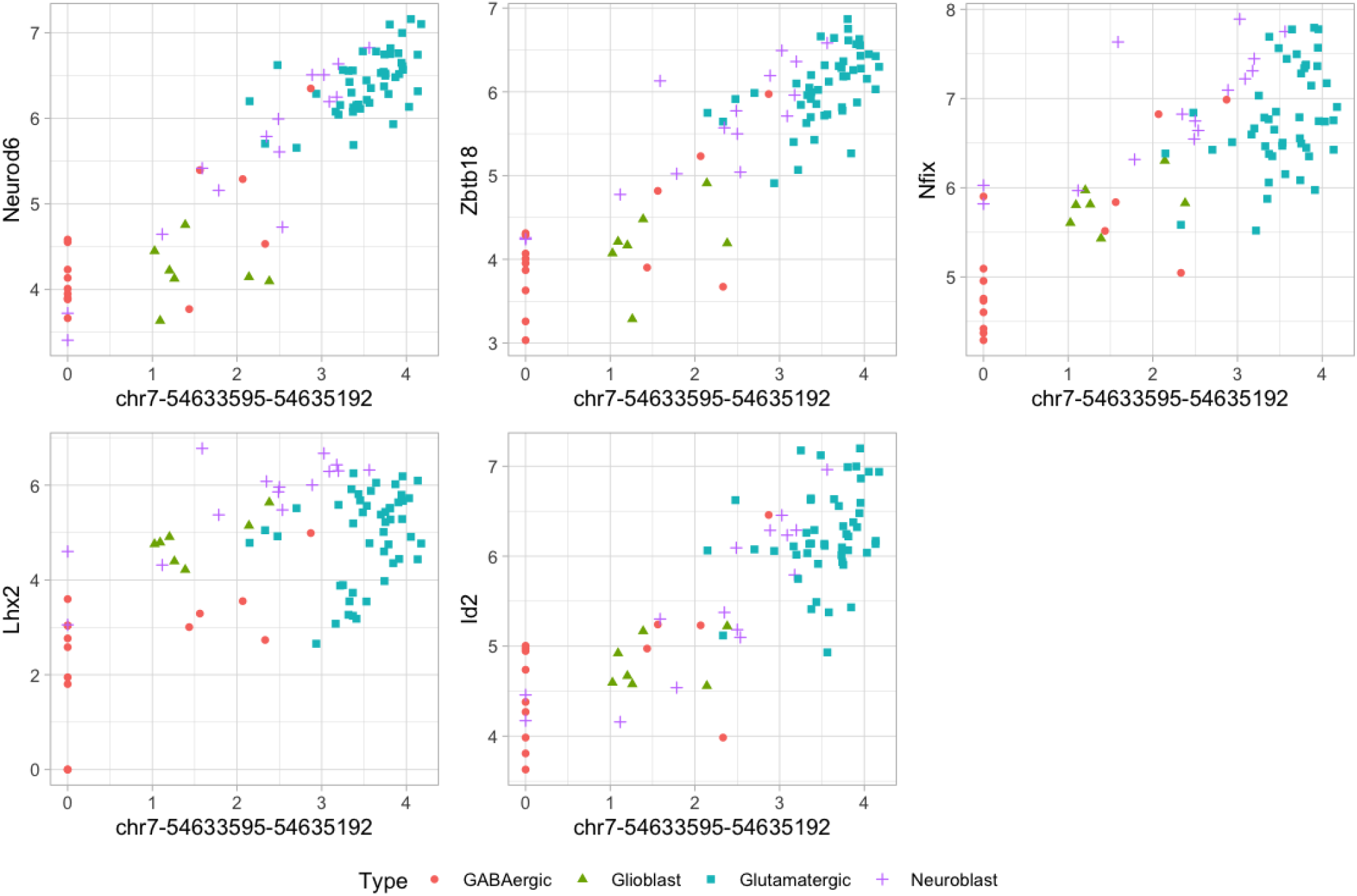
Scatter plots of chromatin accessibility at chr7:54,633,595-54,635,192 and the expression levels of its positively associated genes.

To validate our findings, we compared significant gene-peak pairs with previously reported chromatin interactions, using PLAC-Seq data of the mouse fetal forebrain [Zhu et al., 2019]. Our analysis identified nine significant gene-peak pairs in PLAC-Seq data, all exhibiting positive correlations between chromatin accessibility and gene expression, predominantly in GABAergic and glutamatergic neurons (Figure 8). We also performed *PETScan* to evaluate cis-associations for all genes and their corresponding peaks within 1,000 kb of the transcription start site, identifying 4,211 significant gene-peak pairs with Benjamini & Hochberg (BH) adjusted p-values below 0.05, of which 756 (18.0%) were validated by PLAC-Seq data. These results showed the biological relevance of our approach in capturing chromatin interactions that regulate gene expression in mouse brain development.

**Figure 8:**
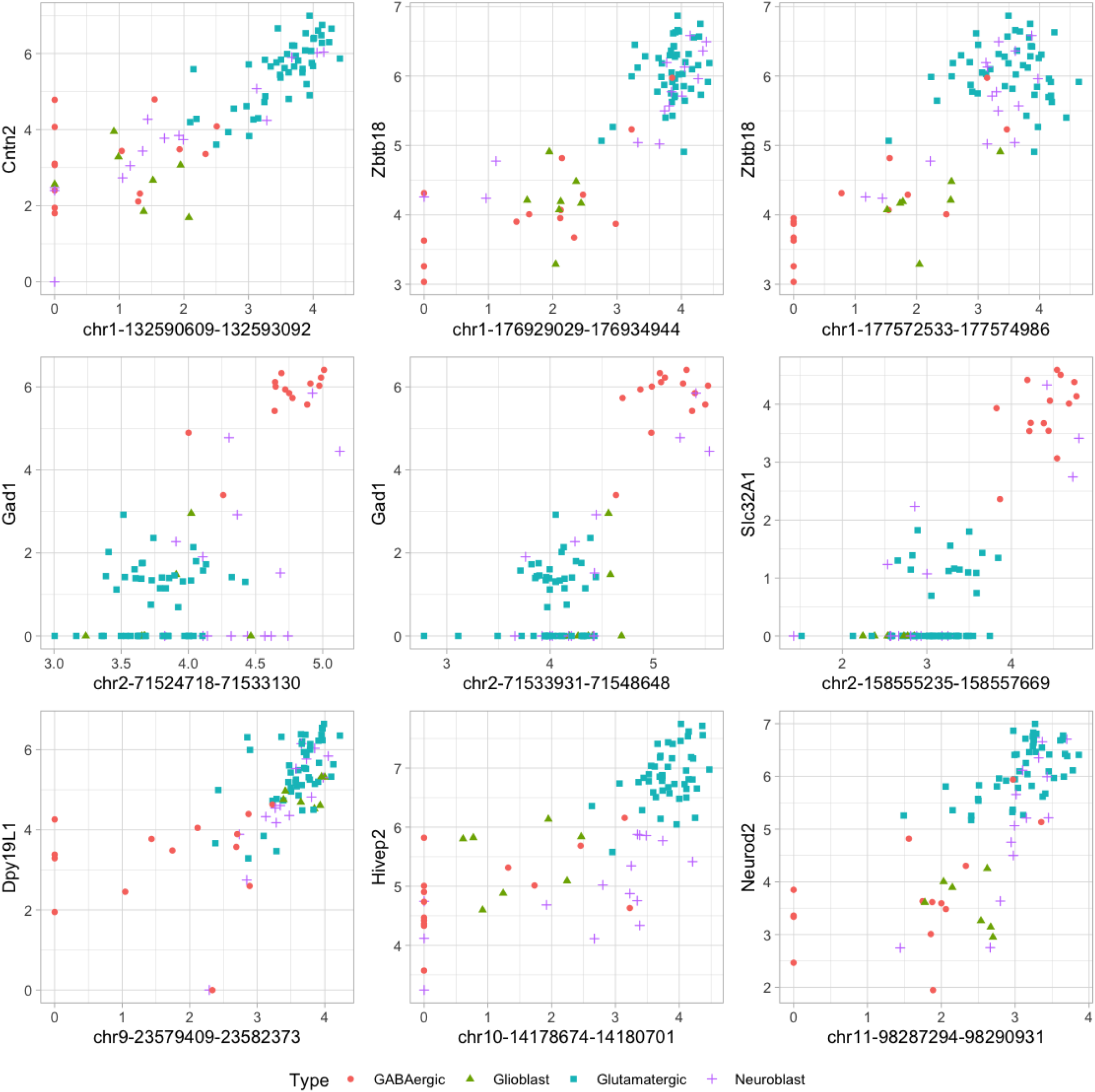
Scatter plots of significant gene-peak pairs from 10x Genomics embryonic mouse brain data validated by PLAC-Seq data.

## 4 Discussion

*PETScan* is a computationally efficient tool for providing a comprehensive perspective on tissue- or cell-type-specific gene regulation and its implications for diverse biological processes and diseases. Our findings align with previously reported cis-eQTLs or PLAC-Seq data, validating the robustness and biological relevance of the identified gene-peak pairs. This concordance underscores *PETScan*’s potential to effectively bridge chromatin accessibility and gene expression data.

Beyond cis-associations, *PETScan* identifies trans-associations, capturing interactions between regulatory elements and distal target genes to uncover long-range regulatory mechanisms that may not be detectable using traditional approaches. Genome-wide associations identified by *PETScan* offer important insights into regulatory mechanisms, providing experimental validation guidelines for confirming the functional roles of specific open chromatin regions in regulating the transcriptomic expressions of their targeted genes.

Although *PETScan* is developed for performing billions of association tests between high-dimensional RNA-Seq and ATAC-Seq data, it is also versatile for other paired omics data, provided that negative binomial models offer an appropriate framework. For example, *PETScan* can be a viable approach for eQTL analysis when gene expression profiles are measured using RNA-Seq data, where the normality assumption of Matrix eQTL is no longer appropriate. Additionally, *PETScan* can facilitate genome-wide association analysis between RNA-Seq and ChIP-Seq data, evaluating how transcription factor binding or histone modifications impact gene expression. Similarly, analysis of paired RNA-Seq and Bisulfite-Seq data with *PETScan* can provide insights into how DNA methylation regulates gene expression.

Less obviously, *PETScan* can be adopted for differential gene expression analysis of RNA-Seq with a given factor, such as a binary outcome. When the sample size is limited, necessitating a large number of permutations, say *P*, to ensure valid type I error control, the *P* sets of permuted outcomes can be treated analogously to chromatin accessibility measurements across *P* peaks, on which *PETScan* can be applied to derive an empirical null distribution. Genomic control can similarly be implemented through permutations to address potential limitations associated with small sample sizes, thereby enhancing the reliability and robustness of the association findings. Similar research direction can be found in [Barry et al., 2025].

A notable feature of *PETScan* is its reliance on score tests, which focus solely on assessing the presence of associations without estimating effect sizes and directionality (positive or negative) as traditional regression-based methods do. While this approach offers computational efficiency and stability, it limits the ability to determine whether a peak positively or negatively influences gene expression, a crucial factor in elucidating underlying biological mechanisms. Nevertheless, *PETScan* remains a powerful tool for large-scale screening in genome-wide association analysis, particularly when computational efficiency is paramount. By identifying candidate interactions for further validation, *PETScan* provides a foundation for subsequent in-depth investigations.

The use of negative binomial models provides the advantage of flexible covariate adjustment compared to simple correlations. Currently, *PETScan* applies the same set of covariates to all genes. However, some covariates may influence the relationship between chromatin accessibility and gene expression in a gene-specific manner, such as the cis-effect of DNA methylation and transcript factor expression levels. *PETScan* can be easily modified to include gene-specific covariates when relevant data is available, enabling more accurate inference.

When multiple paired RNA-Seq and ATAC-Seq samples are collected from the same subject, such as from different biological time points, the correlations between these samples must be accounted for. Our future plan is to extend *PETScan* from negative binomial models to negative binomial mixed models [Tsonaka and Spitali, 2021, Sun et al., 2016, Rivellese et al., 2022], presenting an even more computationally daunting task. In addition, negative binomial models in *PETScan* may not adequately account for counts with excess zeros commonly observed in other omics data, such as single-cell RNA-Seq data and microbiome data, suggesting the need for zero-inflated Poisson or zero-inflated negative binomial models, which could be another useful extension direction for *PETScan*.

In conclusion, *PETScan* offers a computationally efficient and biologically meaningful framework for assessing genome-wide associations between chromatin accessibility and gene expression. It holds promise for advancing the study of gene regulation and guiding experimental validation efforts, ultimately contributing to a deeper understanding of the regulatory mechanisms underlying gene expression.

